# Annexin A4 trimers are recruited by high membrane curvatures in Giant Plasma Membrane Vesicles

**DOI:** 10.1101/2020.02.20.957183

**Authors:** Christoffer Florentsen, Alexander Kamp-Sonne, Guillermo Moreno-Pescador, Weria Pezeshkian, Ali Asghar Hakami Zanjani, Himanshu Khandelia, Jesper Nylandsted, Poul Martin Bendix

## Abstract

The plasma membrane of eukaryotic cells consists of a crowded environment comprised of a high diversity of proteins in a complex lipid matrix. The lateral organization of membrane proteins in the plasma membrane (PM) is closely correlated with biological functions such as endocytosis, membrane budding and other processes which involve protein mediated shaping of the membrane into highly curved structures. Annexin A4 (ANXA4) is a prominent player in a number of biological functions including plasma membrane repair. Its binding to membranes is activated by Ca^2+^ influx and it is therefore rapidly recruited to the cell surface near rupture sites where Ca^2+^ influx takes place. However, the free edges near rupture sites can easily bend into complex curvatures and hence may accelerate recruitment of curvature sensing proteins to facilitate rapid membrane repair. To analyze the curvature sensing behavior of curvature inducing proteins in crowded membranes, we quantifify the affinity of ANXA4 monomers and trimers for high membrane curvatures by extracting membrane nanotubes from giant plasma membrane vesicles (GPMVs). ANXA4 is found to be a sensor of negative membrane curvatures. Multiscale simulations furthermore predicted that ANXA4 trimers generate membrane curvature upon binding and have an affinity for highly curved membrane regions only within a well defined membrane curvature window. Our results indicate that curvature sensing and mobility of ANXA4 depend on the trimer structure of ANXA4 which could provide new biophysical insight into the role of ANXA4 in membrane repair and other biological processes.

## Introduction

Annexin proteins have a number of important functions in cells including maintaining the integrity of membranes. The plasma membrane (PM) is comprised of a complex system of different lipids and proteins including several annexins, which bind peripherally to the inner leaflet in presence of Ca^2+^. The interaction of the annexins with the anionic headgroups in the membrane is important for a number of biological processes including membrane repair, vesicle transport, endo- and exocytosis, membrane trafficking and calcium channel activity.^1^ The protein core domain consist of four conserved homologous repeats (eight in ANXA6) that together form the shape of a slightly bent disc. The Ca^2+^ dependent membrane binding sites are found at the core region on the convex side of the protein, whereas the N-terminal region, which has a large influence on protein stability and function, is found on the concave side. The vertebrate annexin protein family consists of a group of 12 proteins (ANXA1-ANXA11 and ANXA13) and each annexin family member translocates to the PM at distinct levels of Ca^2+^ concentration, thus the annexin family is a broad system of Ca^2+^ sensitive proteins which can be differentially recruited to the a site of injury depending on the local level of Ca^2+^.^2–6^

An important function of the PM is to maintain an osmotic gradient and regulate the intracellular calcium level which is essential for cellular functions. Small ruptures in the PM due to external stress result in influx of Ca^2+^, but cells have developed an efficient PM repair system to cope with membrane ruptures, which relies on several types of calcium sensitive proteins. Annexins have recently received significant attention due to their prominent role in plasma membrane repair. ^2,6^ Proteins from the annexin familiy and in particular ANXA4 have been shown to bind to the area surrounding membrane holes, although their exact role in the membrane repair mechanism remains elusive. ^7^ ANXA4 was recently shown to induce membrane bending or rolling of supported membranes^6,7^ which naturally raises the question whether ANXA4 would sense or induce membrane curvature of the free membrane edges surrounding a plasma membrane rupture.

The interaction between proteins and lipids are diverse and biophysical studies have shown that several classes of proteins can shape membranes or sense membrane curvature. These include Bin/Amphiphysin/RVS (BAR) domains, annexins and epsins.^6,8–12^ Especially BAR domains have been well established as mediators of curvature generation and sensing whereas the curvature sensing abilities of annexins have not been investigated apart from a recent study showing that ANXA5 has a high affinity for the negative membrane curvature inside a membrane nanotube. ^13^ Additionally, ANXA4 and ANXA6 have been reported to generate membrane curvatures when added to supported lipid bilayers.^6,7^ The convex membrane binding site existing in all annexin core domains could be a facilitator for curvature sensing in a similar manner as the convex shaped IBAR domains have been shown to sense negative membrane curvature in membrane nanotubes extracted from synthetic vesicles.^11^ However, despite the similarities between the different annexins, ANXA2 was not found to sense negative membrane curvature in tubes pulled from GPMVs wheres ANXA5 did accumulate inside the same tubes. These differences could be due to the fact that sorting was tested in cellular plasma membranes and protein-protein effects could potentially affect the sorting. Also, it is worth nothing that ANXA5s have been reported to form trimers on membranes whereas ANXA2 has been found to be membrane bound as a monomer. ^14^

From these results, the question arises whether the sorting, observed for some annexins, is dependent on their ability to form multimers. ANXA4 is of great interest as it has been found to trimerize similar to ANXA5.^14–16^ Here, we investigate the curvature sensing of ANXA4 trimers in GPMVs and compare it with an ANXA4 mutant which is unable to oligomerize. Furthermore, we perform parallel multiscale simulations which predict that ANXA4 trimers are indeed both generators and sensors of negative membrane curvature. The simulations further predict that curvature sensing has a non-trivial dependence on the induced local curvature by the protein and that sensing depends on the induced local rigidity increase on the membrane.

## Results and discussion

The complexity of the plasma membrane composition and its interaction with the cytoskeletal filaments makes it difficult to study the interaction of membrane proteins with the membrane *in vivo*. Hence, a more simple system resembling the living cell, and at the same time, containing the factors of interest is desired to eliminate unwanted noise in the experiments. For many years, model systems such as GUVs have been the primary assay for investigating the curvature sensing of proteins and lateral organization of proteins in membranes.^9,11^ Here, we present a way of investigating the membrane curvature affinity of GFP-tagged ANXA4 in a close-to natural environment utilizing GPMVs harvested directly from the cell membrane.

The formation of these vesicles is schematically depicted in Figure 1 together with representative confocal images of the vesicles.

**Figure 1:**
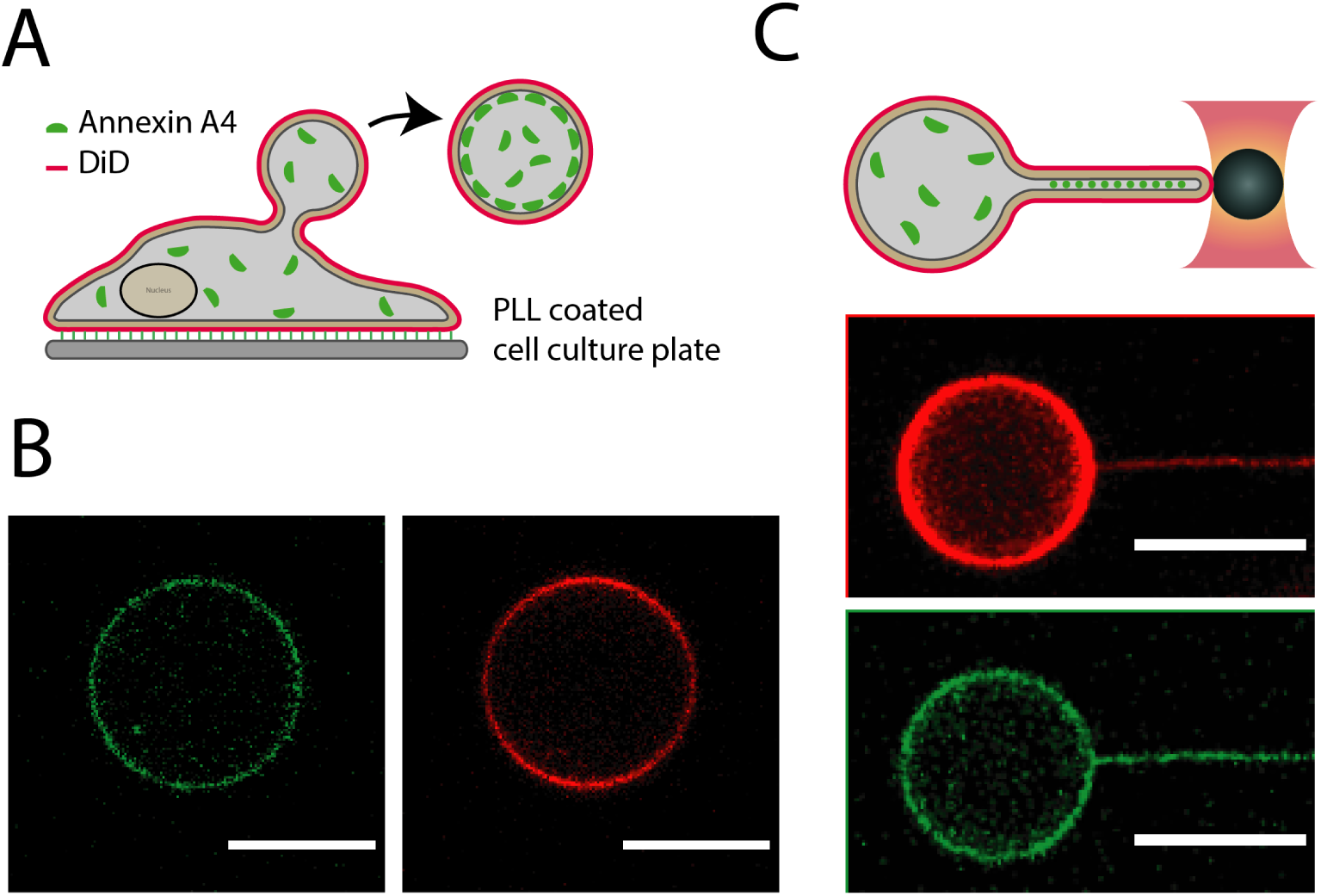
Schematic illustration of the method for producing GPMVs (A). The GPMVs contain ANXA4-GFP which was expressed by the mother cell and leaked into the vesicle during its formation. (B) Confocal images showing representative data of GPMVs with membrane bound ANXA4-GFP (green) and DiD labeled membrane (red). (C) Schematics of experiment where a membrane nanotube is pulled from a GPMV using an optically trapped bead attached to the membrane. Confocal images show the extracted tube from a DiD labeled (red) GPMV containing membrane bound ANXA4-GFP (green), respectively. All scale bars are 10 *µ*m.

To investigate curvature induced sorting and mobility of ANXA4 we use vesicles harvested directly from the PM (Figure 1). The GPMVs are devoid of internal membrane structures like cytoskeleton and nucleus,^17^ but they do contain proteins from the cytosol and transmembrane proteins from the plasma membrane, including proteins produced from transient transfection.^13^ These vesicles are therefore well suited for the purpose of observing protein-lipid interactions in a complex plasma membrane environment. Human embryonic kidney (HEK293T) and breast carcinoma (MCF-7) cell lines were used as donors for the GPMVs and ANXA4-GFP proteins were transiently expressed in these cell lines.

### ANXA4 senses negative membrane curvature

To investigate the curvature affinity of ANXA4, we pulled nanotubes from cell-derived and transfected GPMVs containing ANXA4-GFP. Recruitment of curvature sensing proteins to the highly curved region within the tubes was detected by confocal imaging. Labeling of the GPMV membranes using the lipid analogue DiD, which does not respond to curvature changes in the membrane, ^18^ provided a reference and the DiD signal was compared with the ANXA4-GFP signal to determine if the protein was sorting between the vesicle and the tube.

The sorting of the protein is calculated using equation (1):

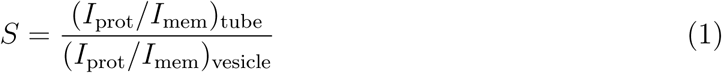

The intensity from the annexin and DiD are noted by *I*_prot_ and *I*_mem_ in eq. (1). The sorting (*S*) denotes the relative density of protein on the tube relative to the flat-membrane reference on the vesicle, *S* = 1 means equal density and no curvature preference whereas e.g. *S* = 5 means five times higher density in the tube.

The sorting values obtained from GPMVs from HEK293T cells transfected with ANXA4-GFP, are shown in Figure 2. There is a clear tendency of ANXA4 to prefer the more curved region inside the nanotube pulled from a GPMV derived from a HEK-293T cell. Almost all measured sorting values are larger than *S* = 1 and range up to 12 times higher density inside the nanotube than on the vesicle membrane. The radii of the nanotubes have not been quantified with a micropipette aspiration experiment and therefore the affinity for specific membrane curvatures was not tested. However, the sorting versus a relative tube diameter is given in Supplementary Figure S1. All the tubes pulled from the GPMVs were below the resolution limit of the microscope (∼ 200 nm) and are expected to be within the range from 20 nm to 200 nm, which has been demonstrated to be the common range of radii for tubes extracted from cells and vesicles.^9,11,18,19^ Occasionally, very thick, abnormal tubes were observed which might be due to contact between the GPMV and a cell-fragment reservoir that is able to add lipids to the system (Supplementary Figure S2). Such tubes were left out of the data shown in Figure 2.

**Figure 2:**
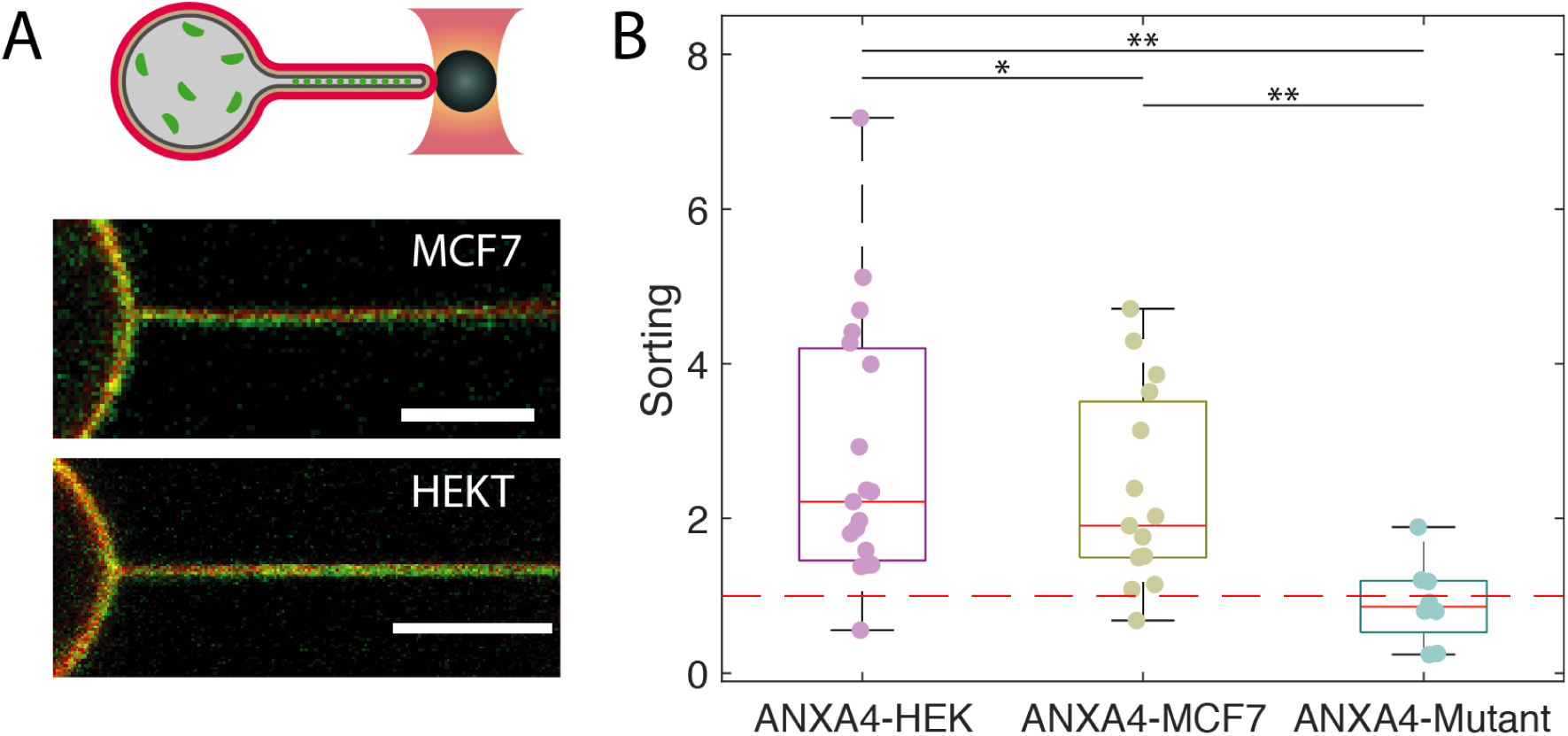
ANXA4 trimers are recruited by high membrane curvatures. A) Membrane curvature sensing by ANXA4-GFP in GPMVs derived from HEK293T and MCF7 cells. Images show overlay of signals from DiD (red) and GFP (green). Scale bars are 5*µ*m. B) The Sorting values indicate the density of protein in the highly curved tube relative to the density on the nearly flat vesicle membrane. The expression of endogenous ANXA4 in MCF7 cells was disrupted by CRISPR editing and expression of ANXA4-GFP was induced by transient transfection thus ensuring presence of only GFP tagged ANXA4 in the vesicles. The ANXA4-trimer mutant was expressed in cells and the GPMVs were used to examine the effect of trimer formation on curvature sensing. The p-values are *p<0.5 and **p<0.01.

To check whether the heterogeneity in the sorting values are caused by a co-existence of transfected ANXA4-GFP and the endogenously expressed fraction of ANXA4, which is not visible in the experiment, we produced GPMVs from a mutant MCF-7 cell line made with the CRISP-Cas9 gene editing technique. In this cell line the production of endogenous ANXA4 was disrupted, thereby eliminating the competition for the binding sites between transiently expressed ANXA4-GFP and endogenous ANXA4 proteins. ^6^ Following transfection with ANXA4-GFP and GPMV formation, nanotubes were pulled and the sorting was quantified in the same manner as for the vesicles from the HEK293T cells. Interestingly, the results show a similar tendency of ANXA4 to migrate to areas of high membrane curvature (see Figure 2), indicating that the curvature sensing properties of ANXA4-GFP are not affected by endogenous ANXA4 proteins.

To determine if the ANXA4 has a preference for positive membrane curvatures, we extracted nanotubes from GUVs and added recombinant ANXA4-GFP to the external medium containing the DiD stained GUVs such that the protein could only bind to the outer surface of the tube. This experiment confirmed that ANXA4-GFP is a sensor for negative membrane curvature which can be concluded from the undetectable protein signal from the nanotube when added to the outside medium, see Supplementary Figure S3. This suggests a lower preference of ANXA4 for positively curved membrane areas than for the “flat” membrane on the vesicle.

The results in Figure 2 show that ANXA4 senses membrane curvatures in MCF7 cells and HEK293T cells in a strikingly similar fashion. The difference observed in the sorting averages between GPMVs from the two cell types, was not significant according to a two sample t-test (p=0.4). These data clearly suggest that having a background of unlabeled ANXA4 in the HEK293T derived GPMVs does not result in different relative sorting when compared with data from MCF7 derived GPMVs which only contained GFP tagged ANXA4. However, there is still a large degree of scattering of data points in both data sets with some tubes exhibiting very low or even negative degree of curvature sorting whereas others are highly enriched with ANXA4. This heterogeneous behavior in sorting could potentially arise from several factors. Firstly, the plasma membrane contains a large number of proteins with a protein area coverage on the membrane that has been reported to reach as high as 30%.^20,21^ This will inevitably lead to crowding effects and possibly other curvature sensing proteins like ANXA5 or IBARs could compete for the same binding sites within the tube. Secondly, the density of ANXA4 on the membrane is not known and could affect sorting as shown for other proteins in artificial systems.^11,22^ Finally, the tubes which were extracted have slightly different radii which could contribute to the heterogeneous data in Figure 2. In Supplementary Figure S1 and S4, we have plotted the sorting data versus relative tube diameter. However, there does not exist a clear inverse correlation between relative tube diameter and sorting, but on average, the sorting is higher for the lower relative tube radii shown in Supplementary Figure S1.

### Multiscale simulations show that ANXA4 trimers generate and sense membrane curvatures

The mechanism behind membrane curvature sensing by ANXA4 could be linked to the convex membrane binding domain of Annexin.

Molecular curvatures have been shown to facilitate curvature sensing and large-scale membrane deformation. ^9–11,23,24^ ANXA4 generates membrane curvature in supported lipid bilayers,^6^ and hence this capacity could also be responsible for the recruitment of the protein to curved membranes. To examine this, we performed a multi-scale simulation by combining all-atom molecular dynamics (MD) and dynamically triangulated Monte Carlo (DTS) simulations. First, we have performed a MD simulation of a system containing one ANXA4 trimer and a lipid bilayer containing POPC and POPS molecules. As expected, the ANXA4 trimer bound to the membrane over the 500 ns timescale and induced negative curvature on the upper bilayer leaflet (Figure 3B). Ca^+2^ ions mediated the interactions of the trimer with negatively charged phosphatidylserine lipids on the bilayer surface. To quantify the induced curvature, we used the last 100 ns of the simulation to find the curvature profile by fitting the phosphorus atom coordinates of the lipids in each monolayer to a Fourier series in two dimensions as it is explained in Ref.^23^ The local mean curvature profile is represented in Figure 3D. The mean curvature under the protein is 0.024 nm^−1^ ± 0.0002 nm^−1^ (corresponding to an induced local curvature of C_0_ = 0.048 nm^−1^ by the ANXA4 trimer).

**Figure 3:**
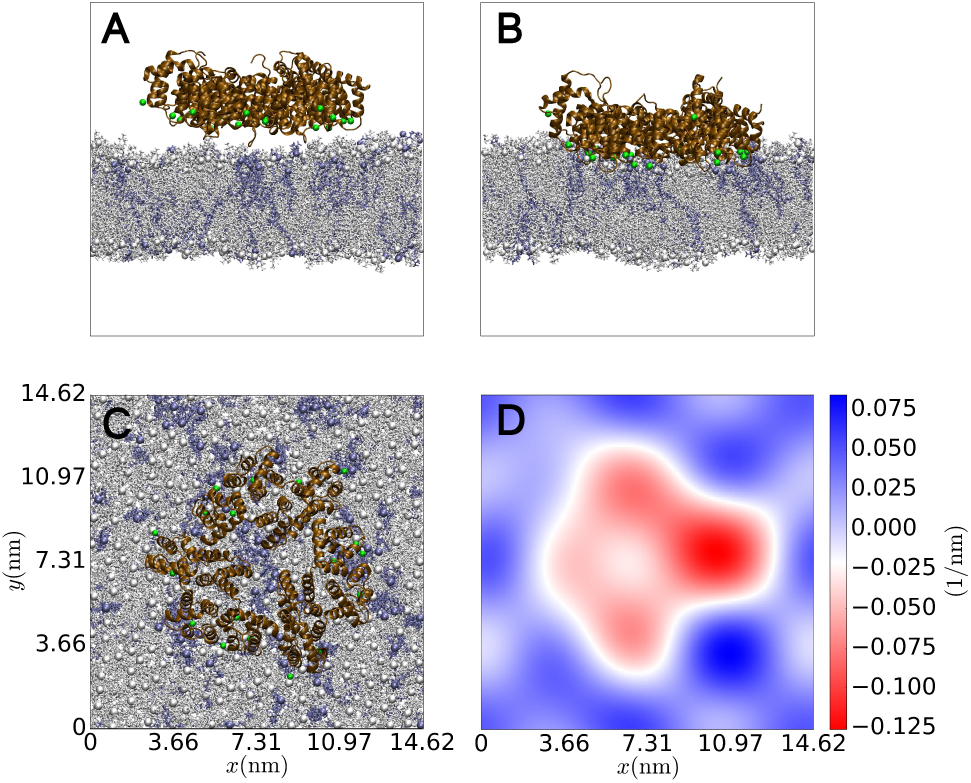
Molecular dynamics simulations showing the membrane curvature imprint imposed by binding of one ANXA4 trimer to a membrane composed of 80% POPC and 20% POPS lipids. POPC lipids are shown in gray and POPS lipids are shown in blue. Ca^2+^ ions are shown as green spheres. (A) lateral view of the initial structure. (B) simulation snapshot of the trimer bound to the membrane showing the indentation of the membrane. (C) simulation snapshot (top view) of the ANXA4 trimer bound to the membrane. (D) The 2D curvature profile for a surface passing through the center of the membrane. The calculated membrane mean curvature is shown by the colorbar and the average mean curvature underneath of ANXA4 trimer is 0.024 ± 0.0002 nm^−1^.

Using the local curvature value and the protein size we calibrate the DTS simulation. Then, we performed DTS simulations of a membrane containing curvature inducing inclusions (protein model).^25^ In our inclusion model, protein-binding modifies local membrane properties by inducing local membrane curvature (a first order coupling to membrane curvature) and local membrane bending modulus (second order coupling to membrane curvature; more details in the method section). ^16^ First, we investigated whether the capacity of ANXA4 to induce curvature can also result in sensing membrane curvature. To do this, we first simulated systems containing 5% annexin coverage for two different values of second order coupling constants (Δ*κ*_*G*_ and Δ*κ*). The results show that (solid lines in Figure 4C, lower panel) annexin effectively senses membrane curvature (higher density on the tube surface) but as (Δ*κ*_*G*_ and Δ*κ*) increases, the sensing capacity decreases. In addition, the results indicate that the presence of the proteins changes the tube diameters in all systems; however, these changes are below the optical microscopy resolution limit to be realized in our experimental setups (Figure 4C, (i) upper panel). Next, we studied the effect of the presence of the other proteins on curvature sensing capacity of annexin. For this purpose, we simulated systems containing 5% annexin, 5% protein with a larger value of induced curvature (C_0_ = 0.062 nm^−1^) than annexin and 5% protein with a lower value of induced curvature (C_0_ = 0.027 nm^−1^) than annexin (Figure 4C, (ii) lower panel). Additionally, we simulated a system containing three proteins all with lower values of induced curvature (Figure 4C, (ii) upper panel). The results show a non-trivial relation between C_0_ and the curvature sensing capacity of the proteins (Figure 4C, (ii) lower panel). First, below a certain threshold of C_0_, proteins do not sense the tube curvature (Figure 4C, (ii) upper panel black curve). In this case the curvature sensing can even become negative which can explain the occasional the negative sensing observed in the experiments. Secondly, above a certain threshold of C_0_, the protein loses its ability to sense curvature (Figure 4C, (ii) lower panel, purple curve). The lower threshold can be understood as a competition between local rigidity induced by proteins (second term in eq. (3)) and the preference of the inclusions for curved regions (first term in eq. (3)). The higher threshold is due to the ability of an inclusion to bend a flat area more easily compared to a tube as the inclusions prefers a cap shape (non-zero local Gaussian curvature) over a tube shape (third term in eq. (3)). Note: this effect cannot be captured by simple theoretical calculations where nanotubes are treated as a perfect cylinder since the Gaussian curvature term is zero on a perfect cylinder.

**Figure 4:**
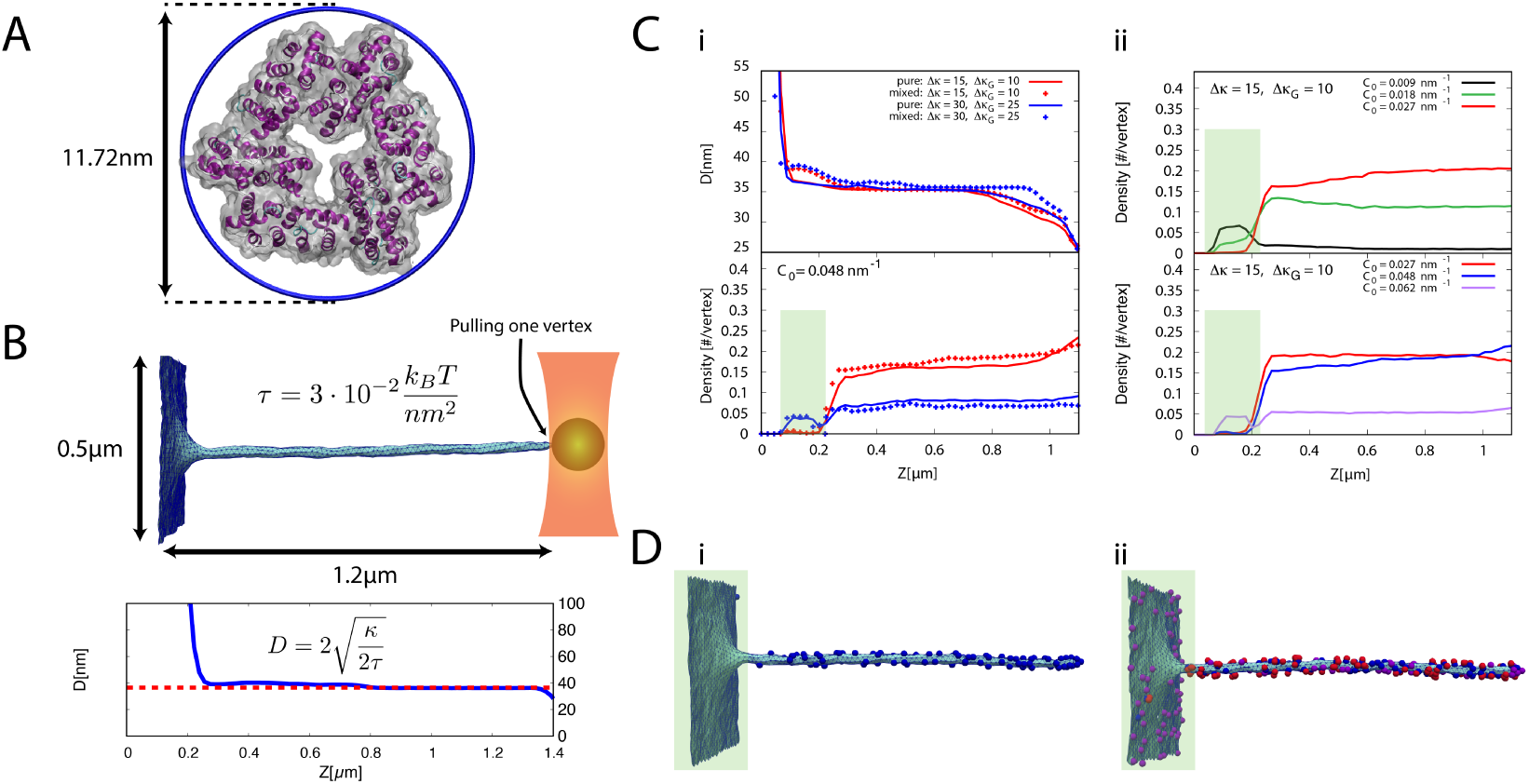
Monte Carlo simulations of ANXA4 protein affinity for a membrane patch (flat) and a highly curved nanotube. (A) structure of the annexin ANXA4 trimer. (B, top) A nanotube was generated by pulling a vertex from a flat membrane patch under tension; *τ* = 4*K*_*B*_*T* (1*/d*^2^), where *d* = 11.2nm is the length metric used in the simulations. (B, bottom) Tube diameter profile corresponding to the image from above, which fits well with theoretical prediction i.e., 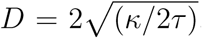, (red dashed line, *κ*_*B*_ = 20*K*_*B*_*T*). (C) (i) Upper panel, tube diameter profile of a tube bound with 5% number density of Annexin4 trimers, for two different values of Δ*κ*_*G*_ and Δ*κ*. (i) Lower panel, Annexin number density profile for two different values of Δ*κ*_*G*_ and Δ*κ*. The solid line is ANXA4 number density profile in absence of any other proteins while the dashed lines are for mixed systems containing 5% number density of other proteins. (ii) Different protein number density profiles in two mixed systems containing different curvature inducing proteins (different *C*_*o*_, for Δ*κ*_*G*_ = 10 and Δ*κ* = 15). Upper panel shows the lower curvature regime and the lower panel shows the higher curvature regime (ANXA4 blue curve). (D) Systems snapshots. Note that the proteins are depicted on the outside of the tube for visibility. (i) System containing only ANXA4 proteins. (ii) System containing ANXA4 and two other different types of curvature inducing proteins (color code is the same as lower panel in (C)(ii)).

These results suggest, for a fixed value of tube radius (in our work 20 nm), only proteins that induce curvature in a small range (in this work 0.01 nm^−1^ *<* C_0_ *<* 0.05 nm^−1^) can sense membrane curvature. The sensing curvature range depends on the tube radius and Δ*κ*_*G*_ and Δ*κ* of the protein model. This can explain why the ANXA 4 sense curvature differently in different system as the tube radius are different in different systems (different membrane bending rigidity and tension that determines the radius of pulled tube). In addition, the results show that the presence of other curvature inducing proteins has a slight effect on the curvature sensing capacity of annexin (comparing solid lines with dashed lines in Figure 4C (i)).

### ANXA4 monomers do not sense membrane curvature

The ability of some annexins to trimerize and form grid-like structures may have important functional importance for their mechanical effect on membranes e.g., during membrane repair.^3,6^ The trimer formation may also affect the proteins ability to sense membrane curvature. In Ref.^13^ it was observed that ANXA5, but not ANXA2, senses negative membrane curvature. Moreover, ANXA5 is known to form trimers in environments with high Ca^2+^ concentration whereas ANXA2 did not show this behavior as reported in Ref.^13^ Since the ANXA4 tested here forms trimers after binding to the membrane^6,26^ we proceeded to investigate the effect of trimerization on the sorting of ANXA4. We introduced a modified version of the mutant protein, engineered not to form trimers (ANXA4-Mutant) in both MCF7 cells and in HEK293T cells. As shown in Figure 2 and Supplementary Figure S4 the trimer-deficient ANXA4-RFP mutant did not show any significant sorting and the difference in sorting for the trimer and trimer mutant was found to be signifcant (p< 0.01) using a two sample t-test, see Figure 2. Hence, these results indicate that the formation of a trimer is essential for the curvature sensing ability of ANXA4.

### Mobility of ANXA4

ANXA4s ability to form trimers could also allow it to form larger connected structures at high protein density as similarily reported for ANXA5 which can form ordered arrays on the PM.^13,15,16,27–29^ ANXA4 has previously been described as being immobile after treating cells with ionomycin, a membrane permeable Ca^2+^ ionophore, and subsequently the lattice formation of ANXA4s can lower mobility of other PM proteins.^29^ From the tubes pulled, we were able to test the ANXA4 mobility using Fluorescence Recovery After Photobleaching (FRAP). By bleaching the nanotubes pulled from the GPMVs, the mobility was monitored as the proteins diffuse inside the tube, see Supplementary Figure S5. However, as seen in Figure 5A, there is not always a protein recovery to detect following bleaching, which is further emphasized by the fact that the nanotube can be pulled further out from the PM vesicle after the FRAP experiment, revealing the former boundary between the vesicle and the nanotube intact (Supplementary Figure S6). This indicates that the ANXA4 is irreversibly bound to the membrane, on the time scale of the experiments, and does not exhibit lateral mobility even at these long time scales. The observation of immobility of the proteins on the PM is further supported by the fact the nanotube stiffness is enhanced as seen from the slow retraction of a nanotube when released from the optical trap (Figure 5B). The relapse of the tube towards the vesicle is not complete when the vesicle contains overexpression of ANXA4-GFP, thus indicating that the ANXA4-GFP changes the physical properties of the tube. This effect was also observed for another curvature generating protein, N-BAR, in Ref.^8^ where the tube force was shown to decrease with the concentration of N-BAR added to the external medium. When tubes were released from GPMVs containing no ANXA4-GFP expression we observed complete retraction of the tubes into the vesicle after ∼ 1s.

**Figure 5:**
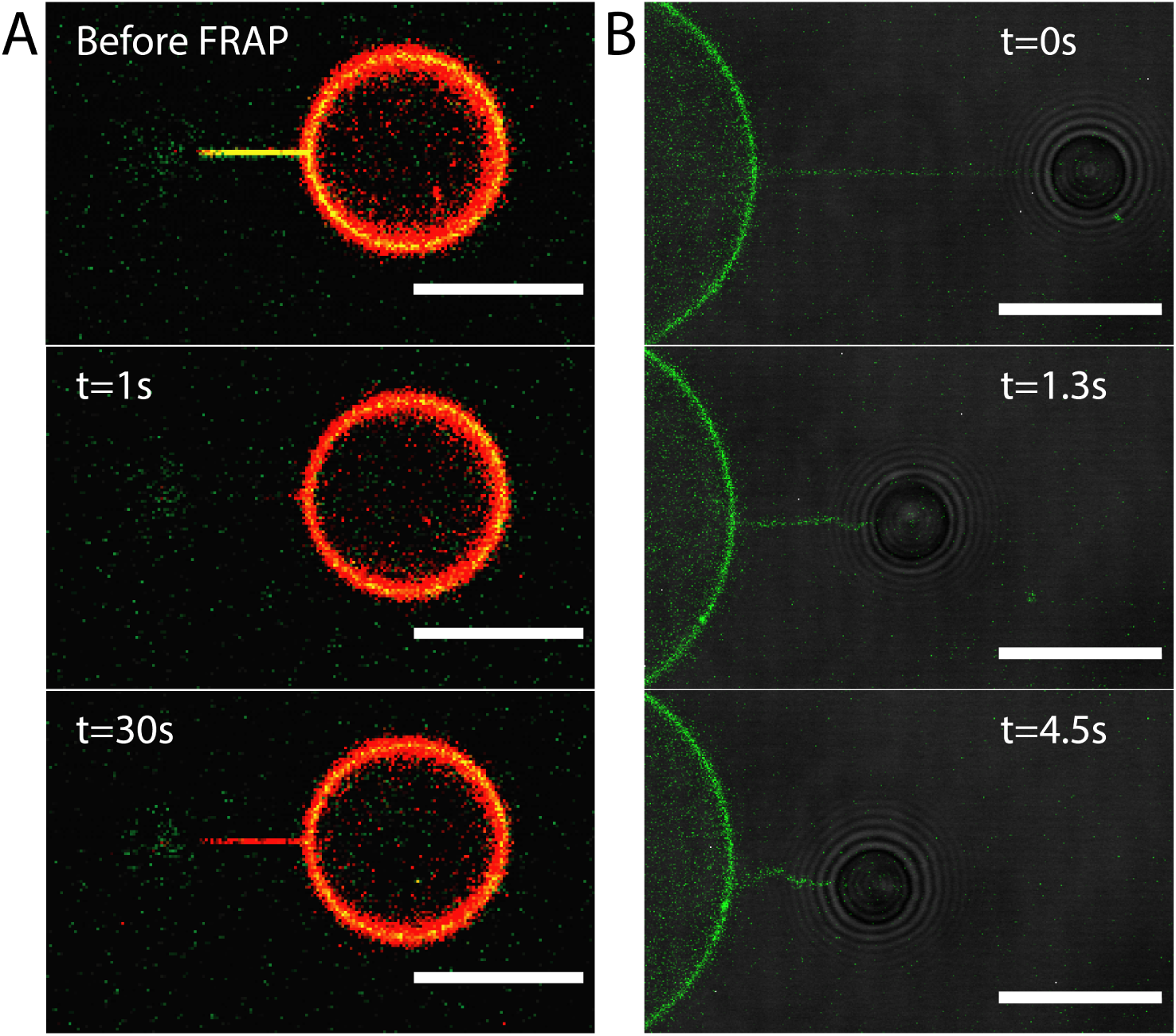
ANXA4 oligomerization stabilizes the nanotube. (A) Overlay images of FRAP experiment showing how the protein (green) is not diffusing into the nanotube after bleaching. After 30s, the vybrant DiD membrane dye (red) is fully recovered, whereas ANXA4 is still not visible in the tube. (B) An example of a nanotube containing ANXA4 (green) which does not become integrated into the vesicle after release from the trap. The retraction of the nanotube is incomplete even after long time periods (>20s), thus suggesting a mechanical effect of the protein on the membrane. Scalebars = 10*µ*m

## Experimental section

### Cell Culture

HEK293T cells (ATCC) were cultured in Dulbecco’s modified Eagle’s medium supplemented with 10% fetal bovine serum (FBS) and 50 units/mL penicillin and 50 *µ*g/mL streptomycin (Fisher). The cells were maintained in a humidified incubator with 5% CO2 at 37°C.

### ANXA4 expression plasmids

Expression plasmids with turbo-GFP C-terminal tag containing human ANXA4, or ANXA5 cDNA were purchased from OriGene Technologies.

ANXA4 mutants deficient in Ca^2+^ binding were generated by PCR-based site-directed mutagenesis by replacing amino acid 71E, 143E (Ca2Mut), or furthermore 227E and 302D (Ca4Mut) in each annexin repeat to A26. To generate an ANXA4 mutant deficient in trimer formation, ANXA5 and ANXA4 alignment was performed to mutate similar amino acids required to inhibit ANXA5 trimer formation (24R, 28-K, 57-K and 192-K to E) (TrimerMut) based on crystal structure evidence16.^13^ ANXA4 mutants were generated in the mammalian pCMV6-AC-mRFP vector to express the proteins with C-terminally tagged red fluorescent protein.^6^

### GPMV Formation

A 6-well plates (Nalge Nunc International, Roskilde, Denmark) was treated with Poly-L-Lysine (PLL) followed by addition of ca. 440000 cells and incubated in appropriate medium over night allowing the cells to adhere properly. When the cells reached 70% confluency, they were transiently transfected (with the plasmid of interest) using Lipofectamine LTX transfection reagent (Fischer Scientific) according to the manufacturer’s protocol. 48 hours post seeding in 6-well plates, the cells PM were stained with DiD (Molecuar Probes, Eugene, OR, USA) followed by washing with GPMV buffer(10 mM HEPES, 150 mM NaCl, 2 mM CaCl2, pH 7.4) three times. For the vesicle formation, the cells were covered with 1 mL of GPMV buffer containing 2 mM NEM and incubated for 1.5-2 hours. Lastly, the vesicles were harvested in the supernatant and left to settle for 30 minutes before use.

### GUV Formation

The composition of the GUV was as follows: 93% DOPC, 5% DOPS and 2 % DiD. The lipids and the dye was mixed at a concentration of 2 *µ*M in chloroform stabilized with ethanol. For the growing buffer, we used 70 mM NaCl (sterile filtered through a 0.2 *µ*m filter), 25 mM Tris (pH 7.4), and 80 mM Sucrose. For the observation buffer, we used 70 mM NaCl (sterile filtered through a 0.2 *µ*m filter), 50 mM Tris (pH 7.4), and 55 mM Glucose.

Polyvinyl alcohol (PVA) gel 5% (w/w) was placed in a 55°C laboratory heating oven for 20 minutes. 90 *µ*L PVA gel was added to plasma cleaned cover glasses and dried at 50°C for 50 minutes. 30 *µ*L of lipid solution was added to the PVA coated cover glass and dried with N_2_ gas followed by drying it in vacuum for 2 hours. The coverglass was placed in a container and protein solution was added followed by addition of the growing buffer (V_total_ = 300*µ*L) and set to form GUVs for 1 hour. The vesicle containing solution was harvested and placed in the observation buffer (1 mL). After 5 minutes, the solution was centrifuged (600rcf for 10 minutes at 13°C) and the pellet isolated and added to fresh observation buffer.

### CRISP-Cas9 edited MCF-7 Cell Line

MCF-7 cells with CRISPR/Cas9-targeted ANXA4 gene disruption is described previously.^6^ Briefly, the following target sequences were used: 5’-GCTTCAGGATTCAATGCCATGG-’3 (#1) using the all-in-one CRISPR/Cas9 plasmid (Sigma-Aldrich) followed by cell-sorting for GFP expression to generate single cell clones with disrupted ANXA4 reading frame.

### Data Analysis

All data analysis was performed using Matlab (The Mathworks, Inc., Natick, MA). Custom-made scripts were used for analysis of the sorting ratios as expressed by eq. 1. Briefly, the vesicle to tube protein ratios were quantified and a membrane dye signal (DiD) was used as reference. Bleaching was accounted for by normalizing to the signal from the vesicle membrane. All Matlab scripts are available upon request.

## Molecular Dynamic Simulation

We built an ANXA4 trimer structure from the ANXA4 monomer (PDB: 1AOW)^24^ by applying the crystal structure information of phosphorylation-mimicking mutant T6D of ANXA4 (PDB: 1I4A)^16^ and using the MakeMultimer.py script.^30^ The membrane is composed of 680 lipid molecules, 20% POPS and 80% POPC, constructed using CHARMM-GUI.^25^ The structure was immersed in a periodic box of TIP3P atomistic water^31^ with 70k water molecules and 1.5 mM salt (405 K^+^, 287 Cl^−^) corresponding to the physiological condition. 18 Ca^+2^ were placed close to some Asp and Glu residues in bottom of the trimer (see Figure 3A). The simulation was performed using Gromacs2019^32^ package and CHARMM36^33–35^ force field. The system was simulated for 500 ns at 310 K using the Nosé-Hoover thermostat. ^36^ The pressure was kept constant at 1 atm using Parrinello-Rahman barostat.^37^

## Monte Carlo simulations

To investigate membrane curvature sensing, using computer simulation, we have employed Monte Carlo (MC) simulation of dynamically triangulated surfaces. In this method, the large-scale conformational properties of a fluid membrane are modelled by a dynamically triangulated, self-avoiding surface subject to periodic boundary conditions in the X and Y directions of the rectangular frame. The Euler characteristics of the surface are zero and therefore the number of vertices *N*_*v*_, triangles *N*_*t*_ and links between vertices *N*_*l*_ are fixed and 6*N*_*v*_ = 3*N*_t_ = 2*N*_*l*_. The self-avoidance is ensured by enforcing the minimum and maximum tether length between neighbouring vertices to be *d* and 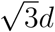 respectively, in which *d* is the length metric of the simulation. Using a set of discretized geometrical operations, a normal vector (N_*v*_), an area (A_*v*_) and two principal curvatures (*C*_1_(*v*),*C*_2_(*v*)) and principal directions (**X**_1_(*v*),**X**_2_(*v*)) are assigned to each vertex.^30^ The bending energy of the system is described by a discretized form of the Helfrich Hamiltonian as

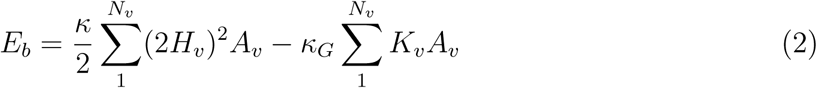

Where *H*_*v*_ = 0.5(*C*_1_(*v*) + *C*_2_(*v*)) and *K*_*v*_ = *C*_1_(*v*)*C*_2_(*v*) are mean and Gaussian curvature respectively and *κ* and *κ*_*G*_ are bending rigidity and Gaussian rigidity. The membrane tension is fixed by a tension controlling algorithm described in.^25^ A protein is modeled as a curvature inducing inclusion that can sit on a vertex and modify its energy by

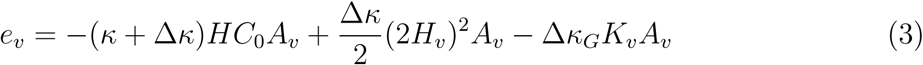

Where C_0_ is the local curvature induced by the protein and Δ*κ* and Δ*κ*_*G*_ represents the local change in the bending and Gaussian rigidities due to the inclusions. The equilibrium properties of the systems are analyzed by Monte Carlo simulation techniques with 4 MC moves i.e., vertex move (a vertex is moved in a random direction), Alexander move (a mutual link between neighboring triangles is flipped and two new triangles will be generated) and Kawazaki moves (an inclusion jumps to a neighboring vertex) and the membrane projected area change. To each Monte Carlo Sweep (MCS) with a probability of *P* = 1*/*(5*N*_*v*_), the membrane projected area is updated and with probability of 1−*P*, trial moves corresponding to *N*_*v*_ vertex moves, *N*_*l*_ Alexander move and *N*_*i*_ Kawazaki moves are performed (for more details see^38–40^). To map the simulation results to the experimental setup, *d*, the simulation length metric, is converted to a physical length. To do this, we assume that the area of a vertex 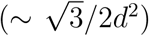 is equal to the area of the smallest circle wrapping an ANXA4 trimer (*A*_*Trimer*_ = 108 nm^2^; Figure 4A) and therefore *d* ≈ 11.2nm.

The nanotube was generated by pulling a vertex with respect to the geometry center of the whole membrane patch using a harmonic potential with a force constant of 60*k*_*B*_*T/d*. The pulling was performed in 10^6^ MCS in which the tube reaches the length of ≈ 104*d* = 1.2*µ*m (Figure 4B). In all simulations, the bending rigidity was set to *κ* = 20*k*_*B*_*T* and after the formation of a nanotube, 10 replica, each for 1.5 · 10^6^ MCS were performed for data analysis (Figure 4C).

The tube radius is calculated as the inverse of the average curvature in a direction in membrane plane and our results show that it matches well with theoretical predictions (Figure 4B, bottom).

### Calculating the tube radius

On each vertex, we find an azimuth tangent direction 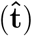 as a unit vector that is parallel to the membrane surface and is perpendicular to the longitudinal axis of the tube (pulling force direction, 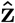). This vector can be found as 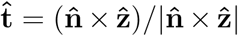, where 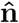 is the normal vector. Then using Euler Curvature formula, we find the surface curvature in the direction of 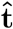 as

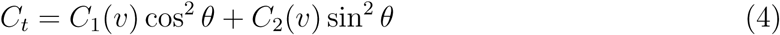

where *θ* is the angle between 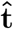 and the direction of the main principal curvature (**X**_1_(*v*)). The tube radius is calculated as inverse of the *C*_*t*_ ensemble average (*R* = 1/⟨*C*_*t*_⟩)

## Conclusion

We have shown that the ANXA4 trimer has the ability to sense negative membrane curvatures and this ability seems to be dependent on trimerization of the protein. The absence of curvature sensing for monomers of ANXA4 is consistent with recent findings that a cognate ANXA2 protein, which exist as monomers on membranes, did not sense membrane curvature. The curvature induced sorting was found to be highly heterogeneous in both plasma membranes harvested from HEK293T cells and from MCF7 cells. Multiscale simulations of protein distributions on a nanotube connected to a flat membrane patch confirmed the ability of ANXA4 trimers to sense membrane curvature. This sensing, however, vanishes when the induced curvature of ANXA4 trimers (*C*_*o*_) is too high or low compared to the membrane curvature. Interestingly, the simulations also predict that ANXA4 has a non-trivial curvature sensing which can change significantly depending on the local increase in rigidity induced by the protein, an effect which cannot be captured by simple theoretical calculations. The the absence of curvature sensing for the ANXA4 monomer remains to be explained and be tested in pure model systems, like GUVs, to see whether the monomer could sense membrane curvature in absence of other proteins. Finally, we observed that ANXA4 could become immobile in the nanotubes which is likely caused by oligomerization of trimers to form a large scale protein lattice. Collectively, these results show that ANXA4 trimers exhibits both curvature sensing and can form a physical scaffold on highly curved membranes which could have important implications in processes such as stabilizing the membrane during plasma membrane repair.

## Acknowledgments

This work is financially supported by Danish Council for Independent Research, Natural Sciences (DFF-4181-00196), by a Novo Nordisk Foundation Interdisciplinary Synergy Program 2018 (NNF18OC0034936), by the Scientific Committee Danish Cancer Society (R90-A5847-14-S2) and by the Lundbeck Foundation (R218-2016-534).

## Supplementary Information

**Fig. S1.**
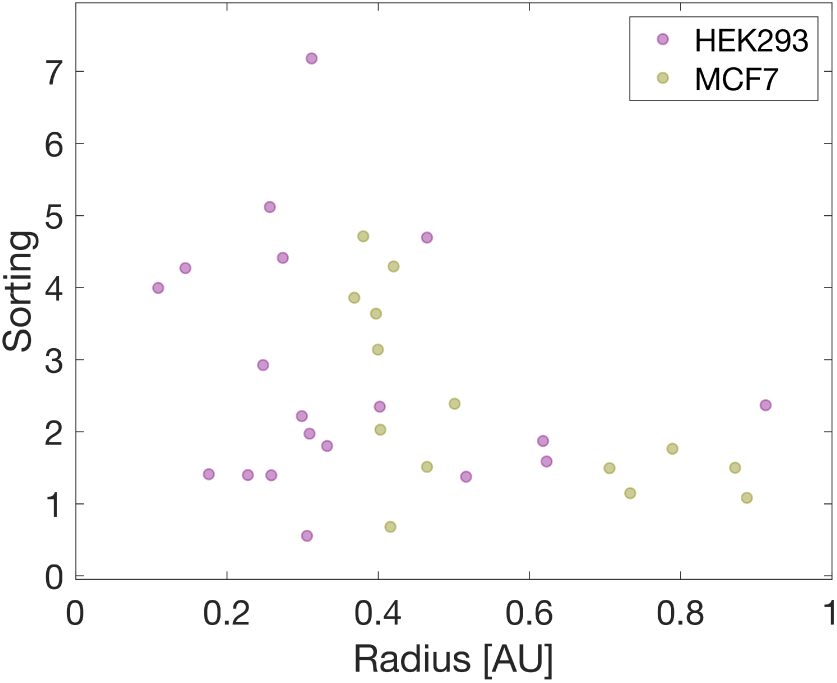
Sorting data as a function of relative tube radius. Data showing the distribution of sorting values for HEK293T cells and MCF-7 cells (unable to produce ANXA4). The x-axis represents the ratio between the DiD intensity measured on the tube and the intensity from the vesicle membrane which scales linearly with the physical tube radius.

**Fig. S2.**
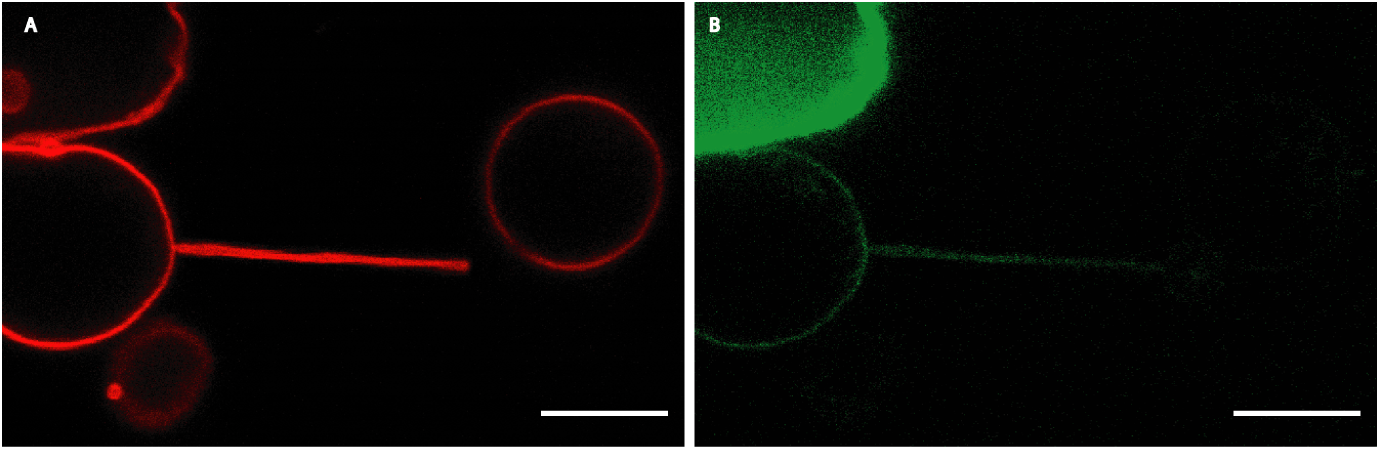
Occasionally, observations of unusually large nanotubes were obtained. The fact that the tether size is so large may stem from the fact that this particular PM vesicle is still attached to a cell fragment which acts as a lipid reservoir, feeding the vesicle lipids and resulting in a low membrane tension. Scale bar = 10*µ*m. A) Shows the DiD channel and B) shows the ANXA4-GFP channel.

**Fig. S3.**
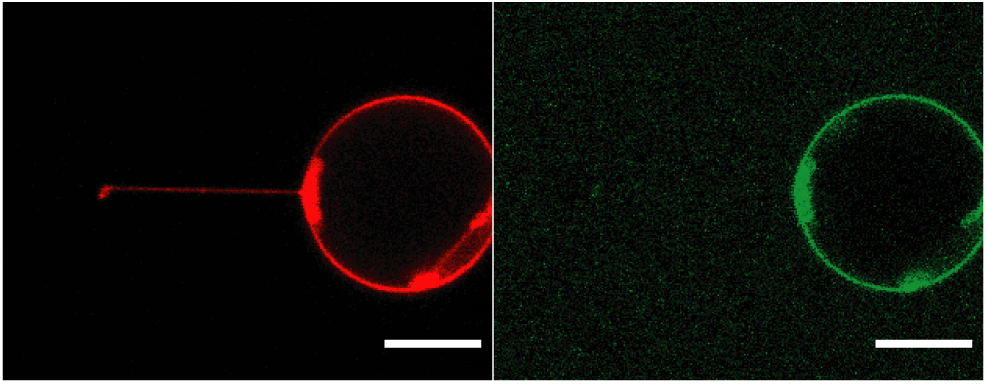
ANXA4-GFP does not sense positive membrane curvatures. Addition of recombinant ANXA4-GFP to the exterior side of GUVs did not result in accumulation of protein on the tube. Left image shows the DiD signal from the tube-vesicle system and the right image shows the corresponding GFP signal from the recombinant ANXA4 protein. Scale bar = 10*µ*m

**Fig. S4.**
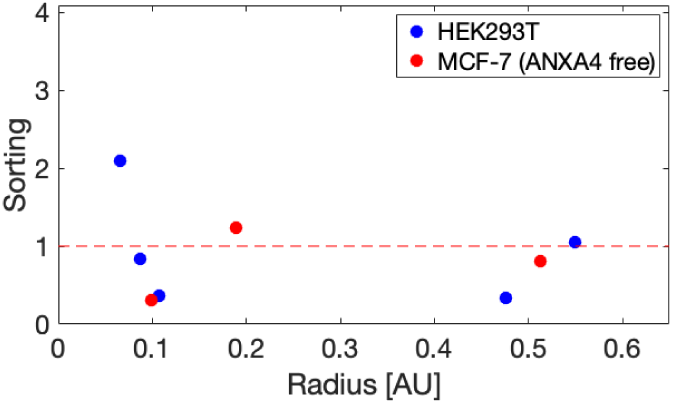
Sorting values for cells expressing the ANXA4-NT-GFP trimer mutant, plotted versus relative tube diameter. The x-axis represents the ratio between the DiD intensity measured on the tube and the intensity from the vesicle membrane which scales linearly with the physical tube radius.

**Fig. S5.**
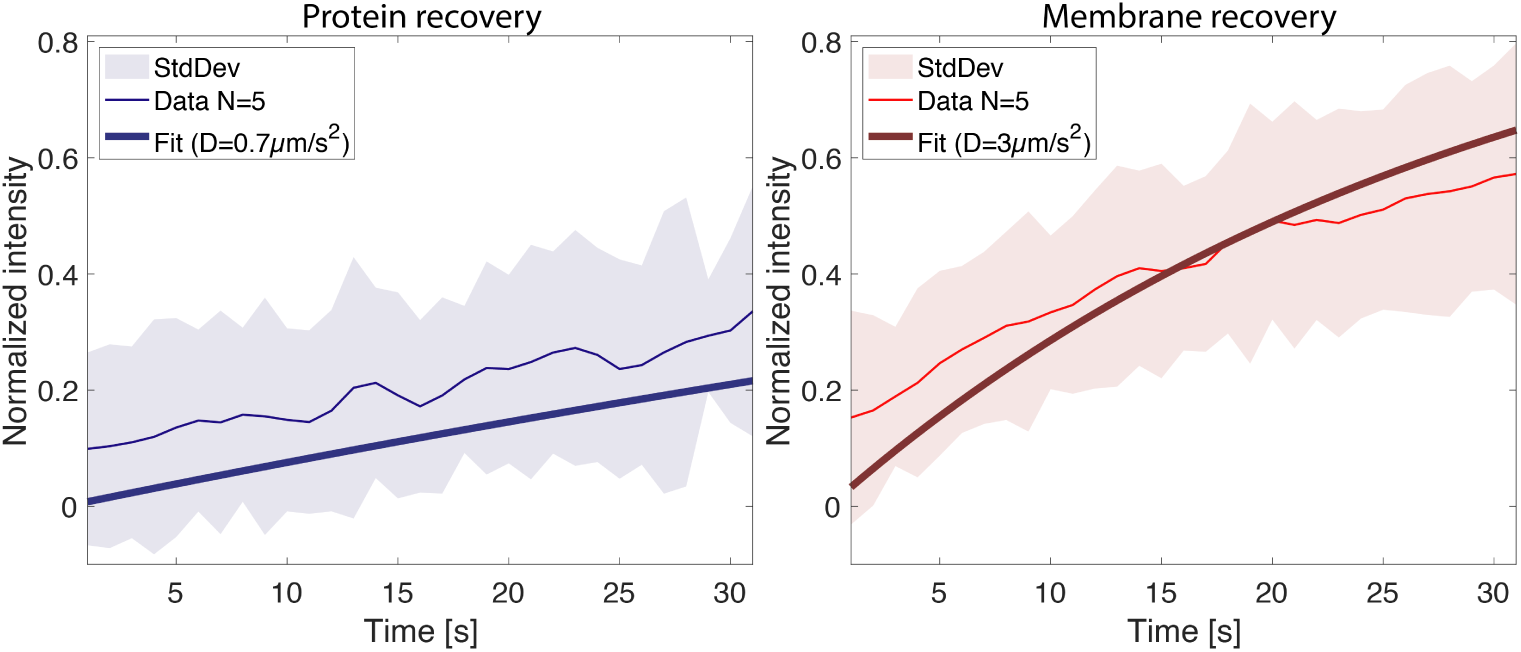
Mobility of ANXA4 measured in experiments where the protein was found to be mobile. Tubes were bleached and recovery curves for 5 experiments were quantified. The diffusion constants were found by fitting the data to the equation: ^13^ 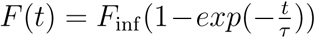 to find the recovery time *τ*. The diffusion constant was then found from 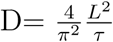 where L is the length of the tube.

**Fig. S6.**
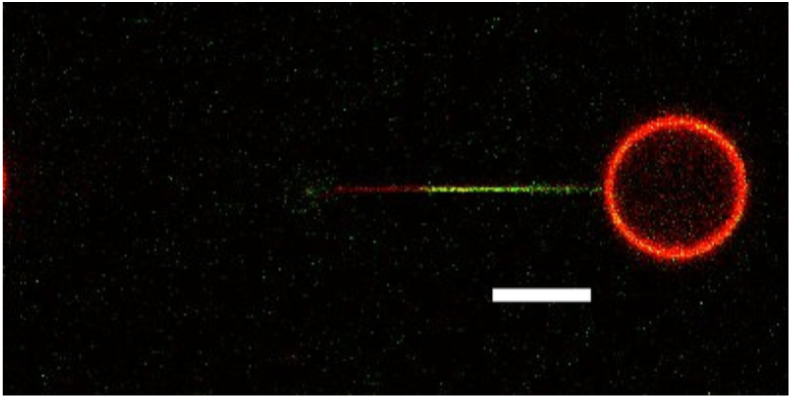
Extension of a nanotube following a FRAP experiment. The immobility of the curvature sensing ANXA4 inside a bleached nanotube is visible even after the nanotube has been extended 20 *µ*m. A clear boundary is visible between the original edge of the nanotube meeting the PM vesicle, and the newly pulled part of the nanotube. Scale bar = 10*µ*m

## References

(1) Lizarbe, M. A.; Barrasa, J. I.; Olmo, N.; Gavilanes, F.; Turnay, J. Annexin-phospholipid interactions. Functional implications. International Journal of Molecular Sciences 2013, 14, 2652–2683.

(2) Draeger, A.; Monastyrskaya, K.; Babiychuk, E. B. Biochemical Pharmacology. Biochemical Pharmacology 2011, 81, 703–712.

(3) Bouter, A.; Gounou, C.; Bérat, R.; Tan, S.; Gallois, B.; Granier, T.; d’Estaintot, B. L.; Pöschl, E.; Brachvogel, B.; Brisson, A. R. Annexin-A5 assembled into two-dimensional arrays promotes cell membrane repair. Nature Communications 2011, 2, 270–9.

(4) Cooper, S. T.; McNeil, P. L. Membrane Repair: Mechanisms and Pathophysiology. Physiological Reviews 2015, 95, 1205–1240.

(5) Lauritzen, S. P.; Boye, T. L.; Nylandsted, J. Annexins are instrumental for efficient plasma membrane repair in cancer cells. Seminars in Cell & Developmental Biology 2015, 45, 32–38.

(6) Boye, T. L.; Maeda, K.; Pezeshkian, W.; Sønder, S. L.; Haeger, S. C.; Gerke, V.; Simonsen, A. C.; Nylandsted, J. Annexin A4 and A6 induce membrane curvature and constriction during cell membrane repair. Nature Communications 2017, 8, 1623.

(7) Boye, T. L.; Jeppesen, J. C.; Maeda, K.; Pezeshkian, W.; Solovyeva, V.; Nylandsted, J.; Simonsen, A. C. Annexins induce curvature on free-edge membranes displaying distinct morphologies. Scientific Reports 2018, 8, 10309–14.

(8) Heinrich, M. C.; Capraro, B. R.; Tian, A.; Isas, J. M.; Langen, R.; Baumgart, T. Quantifying Membrane Curvature Generation of Drosophila Amphiphysin N-BAR Domains. The journal of physical chemistry letters 2010, 1, 3401–3406.

(9) Ramesh, P.; Baroji, Y. F.; Reihani, S. N. S.; Stamou, D.; Oddershede, L. B.; Bendix, P. M. FBAR Syndapin 1 recognizes and stabilizes highly curved tubular membranes in a concentration dependent manner. Scientific Reports 2013, 3, 1565.

(10) McMahon, H. T.; Boucrot, E. Membrane curvature at a glance. Journal of Cell Science 2015, 128, 1065–1070.

(11) Prévost, C.; Zhao, H.; Manzi, J.; Lemichez, E.; Lappalainen, P.; Callan-Jones, A.; Bassereau, P. IRSp53 senses negative membrane curvature and phase separates along membrane tubules. Nature Communications 2015, 6, 8529.

(12) Barooji, Y. F.; Rørvig-Lund, A.; Semsey, S.; Reihani, S. N. S.; Bendix, P. M. Dynamics of membrane nanotubes coated with I-BAR. Scientific Reports 2016, 6, 30054.

(13) Moreno-Pescador, G.; Florentsen, C. D.; Østbye, H.; Sønder, S. L.; Boye, T. L.; Veje, E. L.; Sonne, A. K.; Semsey, S.; Nylandsted, J.; Daniels, R. et al. Curvature- and Phase-Induced Protein Sorting Quantified in Transfected Cell-Derived Giant Vesicles. ACS Nano 2019, 13, 6689–6701.

(14) Patel, D. R.; Isas, J. M.; Ladokhin, A. S.; Jao, C. C.; Kim, Y. E.; Kirsch, T.; Langen, R.; Haigler, H. T. The Conserved Core Domains of Annexins A1, A2, A5, and B12 Can Be Divided into Two Groups with Different Ca2+ Dependent Membrane-Binding Properties. Biochemistry 2005, 44, 2833–2844.

(15) Oling, F.; Bergsma-Schutter, W.; Brisson, A. Trimers, Dimers of Trimers, and Trimers of Trimers Are Common Building Blocks of Annexin A5 Two-Dimensional Crystals. Journal of Structural Biology 2001, 133, 55–63.

(16) Kaetzel, M. A.; Mo, Y. D.; Mealy, T. R.; Campos, B.; Bergsma-Schutter, W.; Brisson, A.; Dedman, J. R.; Seaton, B. A. Phosphorylation mutants elucidate the mechanism of annexin IV-mediated membrane aggregation. Biochemistry 2001, 40, 4192–4199.

(17) Sezgin, E.; Kaiser, H.-j.; Baumgart, T.; Schwille, P.; Simons, K.; Levental, I. Elucidating membrane structure and protein behavior using giant plasma membrane vesicles. Nature Protocols 2012, 7, 1042–1051.

(18) Rosholm, K. R.; Leijnse, N.; Mantsiou, A.; Tkach, V.; Pedersen, S. L.; Wirth, V. F.; Oddershede, L. B.; Jensen, K. J.; Martinez, K. L.; Hatzakis, N. S. et al. Membrane curvature regulates ligand-specific membrane sorting of GPCRs in living cells. Nature Chemical Biology 2017, 13, 724–729.

(19) Tian, A.; Baumgart, T. Sorting of Lipids and Proteins in Membrane Curvature Gradients. Biophysj 2009, 96, 2676–2688.

(20) Dupuy, A. D.; Engelman, D. M. Protein Area Occupancy at the Center of the Red Blood Cell Membrane. Proc. Natl. Acad. Sci. U. S. A 2008, 105, 2848–2852.

(21) Guigas, G.; Weiss, M. Effects of Protein Crowding on Membrane Systems. Biochim. Biophys. Acta, Biomembr. 2016, 1858, 2441–2450.

(22) Sorre, B.; Callan-Jones, A.; Manzi, J.; Goud, B.; Prost, J.; Bassereau, P.; Roux, A. Nature of curvature coupling of amphiphysin with membranes depends on its bound density. Proc. Natl. Acad. Sci. U. S. A 2012, 109, 173–178.

(23) Pezeshkian, W.; Nåbo, L. J.; Ipsen, J. H. Cholera toxin B subunit induces local curvature on lipid bilayers. FEBS open bio 2017, 7, 1638–1645.

(24) Zanotti, G.; Malpeli, G.; Gliubich, F.; Folli, C.; Stoppini, M.; Olivi, L.; Savoia, A.; Berni, R. Structure of the trigonal crystal form of bovine annexin IV. The Biochemical journal 1998, 329 (Pt 1), 101–106.

(25) Lee, J.; Patel, D. S.; Ståhle, J.; Park, S.-J.; Kern, N. R.; Kim, S.; Lee, J.; Cheng, X.; Valvano, M. A.; Holst, O. et al. CHARMM-GUI Membrane Builder for Complex Biological Membrane Simulations with Glycolipids and Lipoglycans. Journal of chemical theory and computation 2019, 15, 775–786.

(26) Kevin C. Crosby, K. C.; Postma, M.; Hink, M. A.; Zeelenberg, C. H. C.; Adjobo-Hermans, M. J. W.; Gadella, T. W. J. Quantitative Analysis of Self-Association and Mobility of Annexin A4 at the Plasma Membrane. Biophysical Journal 2013, 104, 1875–1885.

(27) Miyagi, A.; Chipot, C.; Rangl, M.; Scheuring, S. High-speed atomic force microscopy shows that annexin V stabilizes membranes on the second timescale. Nature nanotechnology 2016, 11, 783–790.

(28) Voges, D.; Berendes, R.; Burger, A.; Demange, P.; Baumeister, W.; Huber, R. Three-dimensional Structure of Membrane-bound Annexin V; 1994.

(29) Piljić, A.; Schultz, C. Annexin A4 self-association modulates general membrane protein mobility in living cells. Molecular Biology of the Cell 2006, 17, 3318–3328.

(30) Palmer, M. MakeMultimer.py.

(31) Jorgensen, W. L.; Chandrasekhar, J.; Madura, J. D.; Impey, R. W.; Klein, M. L. Comparison of simple potential functions for simulating liquid water. The Journal of Chemical Physics 1983, 79, 926–935.

(32) Abraham, M. J.; Murtola, T.; Schulz, R.; Páll, S.; Smith, J. C.; Hess, B.; Lindahl, E. GROMACS: High performance molecular simulations through multi-level parallelism from laptops to supercomputers. SoftwareX 2015, 1-2, 19–25.

(33) Best, R. B.; Zhu, X.; Shim, J.; Lopes, P. E. M.; Mittal, J.; Feig, M.; MacKerell, A. D. Optimization of the Additive CHARMM All-Atom Protein Force Field Targeting Improved Sampling of the Backbone *ϕ, ψ* and Side-Chain *χ*1 and *χ*2 Dihedral Angles. Journal of Chemical Theory and Computation 2012, 8, 3257–3273.

(34) Huang, J.; Rauscher, S.; Nawrocki, G.; Ran, T.; Feig, M.; de Groot, B. L.; Grubmüller, H.; MacKerell, A. D. CHARMM36m: an improved force field for folded and intrinsically disordered proteins. Nature methods 2017, 14, 71–73.

(35) Klauda, J. B.; Venable, R. M.; Freites, J. A.; O’Connor, J. W.; Tobias, D. J.; Mondragon-Ramirez, C.; Vorobyov, I.; MacKerell, A. D.; Pastor, R. W. Update of the CHARMM all-atom additive force field for lipids: validation on six lipid types. The journal of physical chemistry. B 2010, 114, 7830–7843.

(36) Braga, C.; Travis, K. P. A configurational temperature Nosé-Hoover thermostat. The Journal of chemical physics 2005, 123, 134101.

(37) Parrinello, M.; Rahman, A. Polymorphic transitions in single crystals: A new molecular dynamics method. Journal of Applied Physics 1981, 52, 7182–7190.

(38) Pezeshkian, W.; Ipsen, J. H. Fluctuations and conformational stability of a membrane patch with curvature inducing inclusions. Soft matter 2019, 450, 670.

(39) Ramakrishnan, N.; Sunil Kumar, P. B.; Ipsen, J. H. Monte Carlo simulations of fluid vesicles with in-plane orientational ordering. Physical review. E, Statistical, nonlinear, and soft matter physics 2010, 81, 041922.

(40) Pezeshkian, W.; König, M.; Marrink, S. J.; Ipsen, J. H. A Multi-Scale Approach to Membrane Remodeling Processes. Frontiers in molecular biosciences 2019, 6, 59.

